# Spatial patterns of evolutionary diversity in Cactaceae show low ecological representation within protected areas

**DOI:** 10.1101/2022.04.25.489403

**Authors:** Danilo Trabuco Amaral, Isabel A. S. Bonatelli, Monique Romeiro-Brito, Evandro Marsola Moraes, Fernando Faria Franco

## Abstract

Mapping biodiversity patterns across taxa and environments is crucial to address the evolutionary and ecological dimensions of species distribution, suggesting areas of particular importance for conservation purposes. Within Cactaceae, spatial diversity patterns are poorly explored, as well as the abiotic factors that may predict these patterns. We gathered geographic and genetic data from 922 cactus species, which are tightly associated with drylands, to evaluate diversity patterns, such as phylogenetic diversity and endemism, paleo-, neo-, and superendemism, and the environmental predictor variables of such patterns in a global analysis. Hotspot areas of cacti diversity are scattered along the Neotropical and Nearctic regions, mainly in the desertic portion of Mesoamerica, Caribbean Island, and the dry diagonal of South America. The geomorphological features of these regions may create a complexity of areas that work as locally buffered zones over time, which triggers local events of diversification and speciation. Desert and dryland/dry forest areas comprise paleo- and superendemism and may act as both museums and cradles of species, displaying great importance for conservation. Past climates, topography, soil features, and solar irradiance seem to be the main predictors of distinct endemism types. The hotspot areas that encompass a major part of the endemism cells are outside or poorly covered by formal protection units. The current legally protected areas are not able to conserve cactus evolutionary history. Given the rapid anthropogenic disturbance, efforts must be reinforced to monitor biodiversity and the environment and to define/plan current and new protected areas.

## 1. Introduction

A meaningful observation regarding biodiversity is that organisms have uneven distribution across the globe, which can reveal how speciation, extinction, and dispersal events may have impacted species distribution (Lomolino et al., 2009). Naturalists have long been interested in explaining why some regions are biologically richer than others as a way to minimize the Wallacean shortfall of biodiversity knowledge. Mapping biodiversity patterns across taxa and environmental conditions is crucial to address the evolutionary and ecological dimensions of species distribution, such as endemism patterns that emerge in regions with significant concentrations of organisms with little representation elsewhere. Areas such as this are particularly meaningful for conservation purposes (Willing et al., 2003; Graham and Fine, 2008; Swenson et al., 2012; Rosauer and Jetz, 2015).

Endemism has multiple spatial and temporal dimensions that can be related to different taxonomic levels, from families to subspecies (Morrone, 2008). Species-rich areas tend to have high endemism, which is usually correlated with contemporary and historical climate regimes and topography (Sandel et al., 2011; Daru et al., 2015; Barrat et al., 2017; Fenker et al., 2020). The metrics used to estimate endemism (see appendix A in supplementary material) are influenced by the defined spatial scale, which can range from large (e.g., continent) to small areas (mountain tops). On a temporal scale, endemism can be described as a result of recent speciation with no dispersion out of the ancestral area (neoendemism) or as the persistence of lineages extinct elsewhere (paleoendemism) (Stebbins and Major, 1965). Such patterns could also be associated with the concepts of cradles (neoendemism) and museums of biodiversity (paleoendemism) and they are not mutually exclusive in geographical space (mixed-endemism; Mittelbach et al., 2007).

While traditional approaches to capture endemism rely on taxonomic measures such as the number of endemic or range-restricted taxa in a region (Kier et al., 2009), approaches using phylogenetic endemism (PE) are more inclusive by accounting for the evolutionary history underlying endemism inferences (Mishler et al., 2014; Sander et al., 2020). To integrate the biological diversity and phylogenetic singularity, the PE metric weights the branch lengths of each lineage by their respective geographic ranges (Rosauer et al., 2009). For example, the observation of long branches restricted to a small geographic area is interpreted as high PE. Under this approach, endemism is not focused exclusively at the species level, as clades at all levels can also be endemic, encompassing intra- to interspecific scales (Mishler et al., 2014). As a consequence, PE patterns can be better related to evolutionary processes and biogeographic events responsible for the changes in speciation and extinction rates (Davies and Buckley, 2011; Schluter and Pennell, 2017).

Areas with historical climatic instability tend to harbor fewer endemic species, often represented by phylogenetically derived species (neoendemics) (Jansson 2003, Sandel et al., 2011). Conversely, historical stable areas may have allowed the survival of ancient lineages, which have been extinct elsewhere (paleoendemics) (Fjeldsa et al., 1999). As a result, paleoendemisms and neoendemisms have different effects on PE inferences. The local extinction of paleoendemic lineages, for instance, increases patterns of PE (Daru et al., 2020). However, the loss of neoendemic lineages would strongly impact the PE only if the entire clade disappeared. Dispersal rates are also important influences on PE. While higher dispersal rates reduce the concentration of endemic species, poor dispersal ability increases endemism (Daru et al., 2017).

Contrasting biodiversity patterns based on phylogenetic information is of special interest in systems in which taxonomy is in flux and may not reflect true lineage diversity, such as the Cactaceae family. These plants have an endemic distribution covering the Americas (except for *Rhipsalis baccifera*) and are associated with a myriad of xeric environments, soil textures, solar irradiance, and altitude (Parker, 1988; Taylor and Zappi, 2004). Cactaceae is a charismatic group of plants and a bona fide example of recent radiation (Arakaki et al., 2011), harboring remarkable diversity in growth forms (Hunt et al., 2006). Additional reasons to assess the endemic levels within the family are to evaluate the levels of endemism, the number of species with disjunct and small range sizes, the threatened IUCN criteria for many species, and the insufficiently protected areas for the group (Goettsch et al., 2019). Moreover, the low level of protected areas may cover just a small portion of the cactus occurrence, which needs to be evaluated and formally recognized in semiarid and arid lands in the Neotropics.

Here, we explored geographic and genetic data from cactus species to conduct an evaluation of the diversity pattern for this plant family. We addressed three main questions: (1) compare spatial patterns of species distribution, (2) identify putative drivers of the spatial patterns of endemism, and (3) evaluate gaps for species protection based on the distribution of protected areas. We hypothesize that long-term stable areas such as historical refugia concentrate cactus evolutionary diversity and show high levels of PE. In addition, based on the taxon requirements, we hypothesize that the main abiotic factors driving the diversity patterns in the family are climate conditions (temperature and precipitation) and environmental heterogeneity which affects species persistence and dispersal along with geographic space (see appendix B in supplementary material for more details about the hypotheses tested).

## 2. Methods

### 2.1. Species distribution data

The Cactaceae species found in the Nearctic and Neotropical regions, extending from southern Canada to southern South America, were used in this study. The species *Rhipsalis baccifera*, the only naturally present in and outside of the American border, was subsampled to the Americas. We used three geographic databases to recover species occurrences, the Global Biodiversity Information Facility (GBIF), iNat (iNaturalist), and iDigBio (Integrated Digitized Biocollections), using the *spocc* package in R (Chamberlain et al., 2021; R Core Team, 2019) between March 10th and 12th, 2021. The Caryophyllales.org checklist (Korotkova et al., 2021) was checked to remove any taxa with incorrect names or misspellings. Points with spatial uncertainty, inappropriate, or inaccurate localization were removed using the *CoordinateCleaner* package in R (Zizka et al., 2020).

### 2.2. Molecular data and phylogenetic analysis

The genetic data were recovered using the *phytotaR* package in R (Bennett, et al., 2018) and the script available at the phylotaR GitHub page (https://github.com/ropensci/phylotaR/). The genes were aligned using MAFFT v.7.31 (Katoh et al., 2002) and concatenated to generate a supergene using the program *catfastat2phyml.pl* (available at: https://github.com/nylander/catfasta2phyml). The topology reconstruction was carried out in IQTree v.2.1 software (Nguyen et al., 2015) using 10,000 ultrafast bootstrap replicates, contracting branches with less than 50% bootstrap support values using the iTOL v.6.4.2 online web server (Letunic and Bork, 2006; available at: https://itol.embl.de/). Considering the potential bias on datasets, phylogenetic reconstruction of the Cactaceae tree was performed using three datasets with distinct amounts of missing data (MD; 40%, 60%, and 80%; supplementary material). We compared the phylogenetic tree topologies with the symmetric Robinson–Foulds (RF) pairwise distance (Robinson and Foulds, 1981) in the R package phytools (Revel, 2012) and performed a Principal Coordinate Analysis (PCoA) in the R package *PCAmixdata* (Chavent et al., 2017). We also conducted experimental pilots using the three MD datasets and observed a similar diversity pattern, except for the 40% MD dataset, which was an outlier in PCA, showed shorter branches, and displayed the highest level of PD. Thus, we used the tree topology (Fig S2) generated with the dataset containing 80% missing data. After phylogenetic reconstruction, we crossed the information among species with geographical coordinates and the species with genetic information, summarizing the final dataset to 922 species (~53% of the total of described species) from 130 genera (~92% of the total of described genera; Korotkova et al., 2021). This dataset was used in the downstream analyses.

### 2.3. Calculation of diversity metrics

The geographic coordinates and the phylogeny recovered in this study were imported into Biodiverse v.3.1 (Laffan et al., 2010) using the Biodiverse pipeline in R (https://github.com/NunzioKnerr/biodiverse_pipeline). We defined 100 x 100 km grids (1° x 1°), which generated 1,963 grid cells covering the Nearctic and Neotropical regions. We matched our occurrence dataset to the mapped grid cells and incorporated our phylogenetic relationship and branch lengths into the analysis. The metrics calculated were: taxon richness (TR), weighted endemism (WE), phylogenetic diversity, relative phylogenetic diversity (RPD), phylogenetic endemism (PE), and relative phylogenetic endemism (RPE; for more details see appendix A in the supplementary material). The statistical significance of PD, PE, RPD, and RPE of each grid cell was estimated using a null model that randomly reassigns the species to each grid cell. The randomizations were run 999 times and the grid cells were classified as significantly high or low (values higher than 97.5% or lower than 2.5%, respectively).

We also performed endemism categorization using CANAPE (Categorical Analysis of Neo- and Paleo-Endemism) and evaluated statistical significance using randomization-based tests in Biodiverse software. We estimated endemism based on weighted endemism (WE), phylogenetic diversity (PD), relative diversity (RPP), weighted phylogenetic endemism (WPE), and relative endemism (RPE). In order to determine the areas of high diversity patterns within formally protected areas, we also overlapped the endemism, PD, and PE maps with shapes of protected areas in the Neotropical and Nearctic regions (UNEP-WCMC and IUCN, 2021), using QGIS 2.18 software (QGIS Development Team, 2009).

### 2.4. Abiotic correlates of diversity pattern

To assess the correlation among spatial patterns of endemism, PD, PE, paleo-, neo-, mixed, and super endemism obtained in this study with abiotic features, we compiled 49 environmental variables, including current and past climatic, topographic, solar irradiation, and soil features from public databases (see Table S1 for more details). Due to uniformity resolution among variable rasters, we resampled all of them to 50 km resolution using the function *resample* present in the R package *raster* (Hijman, 2019). We also removed the collinear variables using the Variance Inflation Factor (VIF), present in the R package *usdm* (Naimi et al., 2014). In this analysis, we used a threshold of 10, which reduced from 49 to 17 abiotic variables: present and past precipitation (wc_Bio15, wc_Bio18, and wc_Bio19; lig_Bio14, lig_Bio15, lig_Bio18, and lig_Bio19), past temperature (lig_Bio2, lig_Bio3, lig_Bio8, and lig_Bio9), topography (terrain roughness index, current_tri; wetness index, current_topoWet), soil features (sq1-nutrient availability, sq1; oxygen viability to the root, sq4; and texture and phase, sq7), and solar irradiance (direct normal irradiation, dni; see more details in Table S1). To identify possible abiotic variables that may predict the spatial pattern of diversity, we ran, trained, and ensembled the prediction of the correlative model obtained from four machine learning algorithms using a modified version of the script described by Paz et al. (2020).

## 3. Results

The spatial distribution and phylogenetic datasets were applied to investigate the diversity pattern associated with the geographical distribution of the Cactaceae family. The phylogenetic tree topology recovered here (Fig. S2) the main major Cactaceae clade proposed by Guerrero et al. (2019). The relationships between the minor Cactaceae clades were also quite similar, with exceptions made for relationships within some genera of tribe Cacteae, especially considering the taxa from genus *Turbinacarpus* (Fig. S2). It is worth noting the inconsistency in the position of this genus, which was also recovered in previous studies (Vázquez-Sánchez et al. 2013; 2019).

### 3.1. Geographic estimates patterns

Cactaceae has a distribution predominantly associated with open and xeric formations that usually display distinct annual temperatures, solar irradiation, and precipitation levels (Bonatelli et al., 2014; Guerrero et al., 2016; Lavor et al., 2020; Sarmiento, 2021). Furthermore, the diversity pattern is unevenly distributed among these areas (Fig. 1). Here, we observed three main cores of high endemism (from north to south): i.) the Chihuahuan desert + Sierra Madre Oriental; ii.) Dry Chaco + the Sechura Desert/Atacama Desert/Chilean Matorral, and iii.) the South Brazilian Atlantic forest and part of the Espinhaço Range (Brazil), the second largest mountain chain of South America. We also identified small hotspot areas in the savanna patch in the south of Florida and Panama’s central portion, and in the Espinhaço Range (Fig. 1b). The high PE areas also coincided with the regions of weighted endemism, while the estimated areas of phylogenetic diversity (PD) were more striking than the endemism and PE cell grid (Fig 1c-d). High PD areas comprised a large portion of the central-northern region of Mexico (desert and xeric shrubland areas in North America), the Dry Chaco, from the southern part of the Brazilian Atlantic forest to Pampas, and a portion of the Espinhaço Range and southern Andean steppes, both areas of remarkable species richness.

**Fig. 1.**
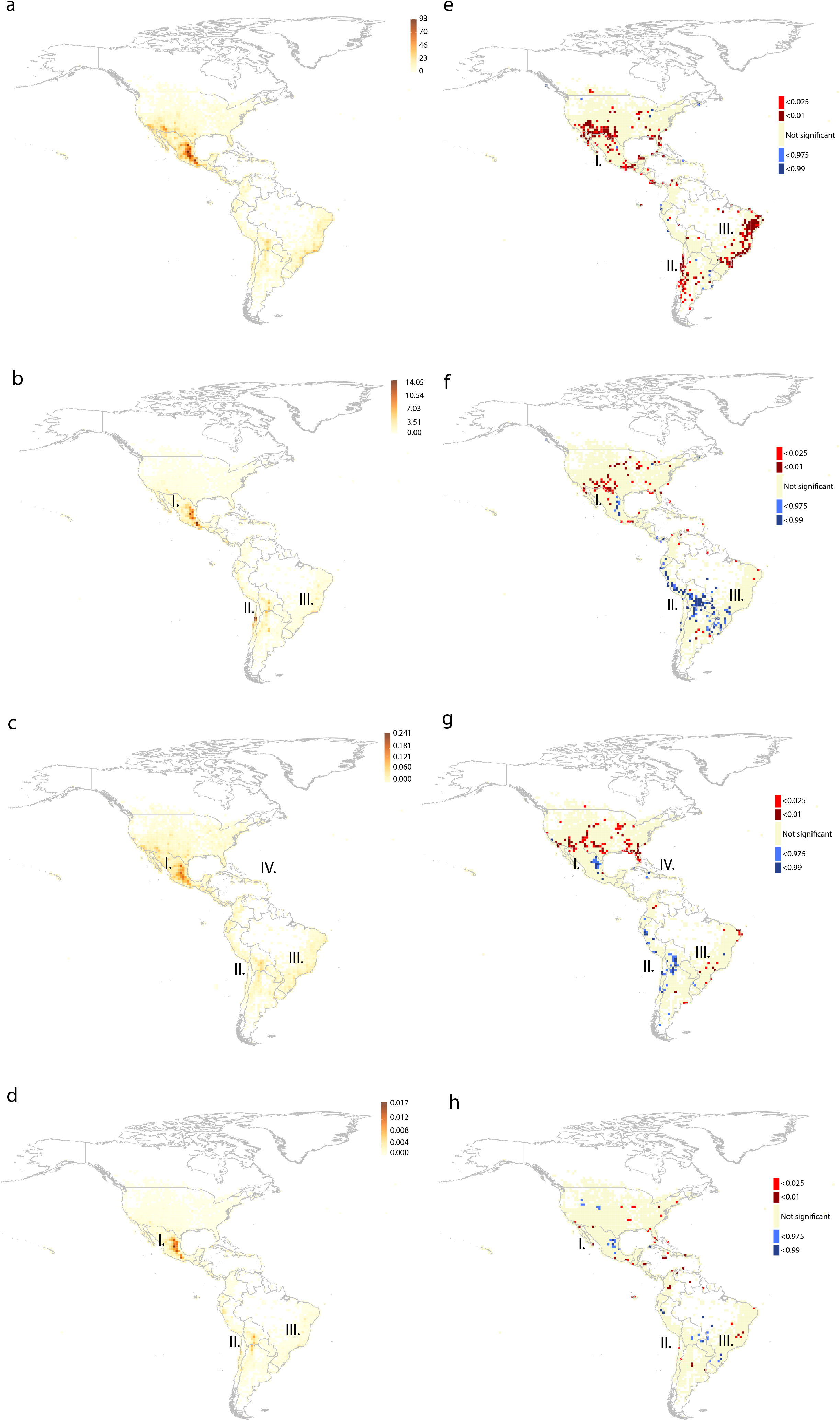
Spatial phylogenetics of the 922 cacti species showed four cores of high diversity in the Neotropical region. a) Taxon richness, (b) weighted endemism (WE), (c) phylogenetic diversity (PD), (d) phylogenetic endemism (PE), (e and f) distribution of phylogenetic diversity (PD), and relative phylogenetic diversity (RPD), and (g–h) similar plots of phylogenetic endemism (PE) and relative phylogenetic endemism (RPE) for Neotropical cacti. Blue and red cells show areas with significantly high (> 0.975) and significantly low (< 0.025) randomized values. Roman numbers represent the four cores of the diversity pattern: I.) the Chihuahuan desert + the Sierra Madre Oriental; II.) Dry Chaco + the Sechura Desert/Atacama Desert/Chilean Matorral, III.) southern Brazilian Atlantic forest and part of the Espinhaço range (South America), and IV.) the Caribbean islands.

There were spatial discrepancies between the PD and relative phylogenetic diversity (RPD), PE, and relative phylogenetic endemism (RPE) patterns (Fig. 1e–h), which were mainly correlated with significantly lower PD values along the Brazilian Atlantic coast. The central portion of Mexico and the Andes/Dry Chaco cells recovered with the highest PE, RPE, and RPD values. Areas with significantly low PD were more common than areas with significantly high PD, recovering only a small portion of the family distribution. These areas were also recovered in both Neotropical and Nearctic realms, mainly along the borders of the Sonoran and Chihuahuan Deserts, Atacama Desert, and Caatinga+Brazilian Atlantic coast. However, the highly significant PD areas involved part of the Chaco, Caribbean islands, and the northeastern portion of the Pacific coast, in Peru and Ecuador. Areas of significantly low and high RPD, PE, and RPE (Fig. 1f-h) showed a similar pattern; three main areas of significantly high values in the central portion of Mexico, into the Andes+Dry Chaco, and South of Brazil, while significantly low values were observed in a large portion of the United States and the northeastern portion of the Atlantic Forest in Brazil.

### 3.2. Paleo-, neo-, and superendemism spatial patterns

We identified five areas of paleoendemism, which included the central-east portion of Mexico, the North Andes portion in Peru, western Cuba, the central portion of Bolivia (Intersection between Dry Chaco and Chiquitania), and southern Brazil (Fig. 2). Seven areas of expanded neoendemism were identified across the Americas, occurring in the savannah portion of eastern USA, south of Baja California, Caribbean islands and south of Florida, south of Mexico, north portion of South America (Venezuela+Guiana), Atacama Desert, Dry Chaco, and southern portion of Espinhaço Range. The expanded mixed endemism, which involved part of Baja California and Mexico, Caribbean islands, eastern Brazil, most of the Andean region and Dry Chaco, and southeastern Argentina, recovered most of the grid cells classified as significantly high PE by CANAPE (Fig. 2). Four main portions of superendemism were identified: in the California Chaparral, the Chihuahuan Desert+Sierra Madre in Mexico, central-eastern Peru, Atacama Desert, Dry Chaco, and southern Espinhaço Range (details in Fig. 2). Expanded areas of mixed and super endemism were commonly detected across the Chihuahuan Desert, Atacama Desert, and Andean regions. It is worth noting that only the southwestern portion of the Dry Diagonal of South America, which comprises the Dry Chaco, displayed significant levels of endemism of any kind, with a low contribution of Brazilian savannah and Caatinga (Fig. 2c).

**Fig. 2.**
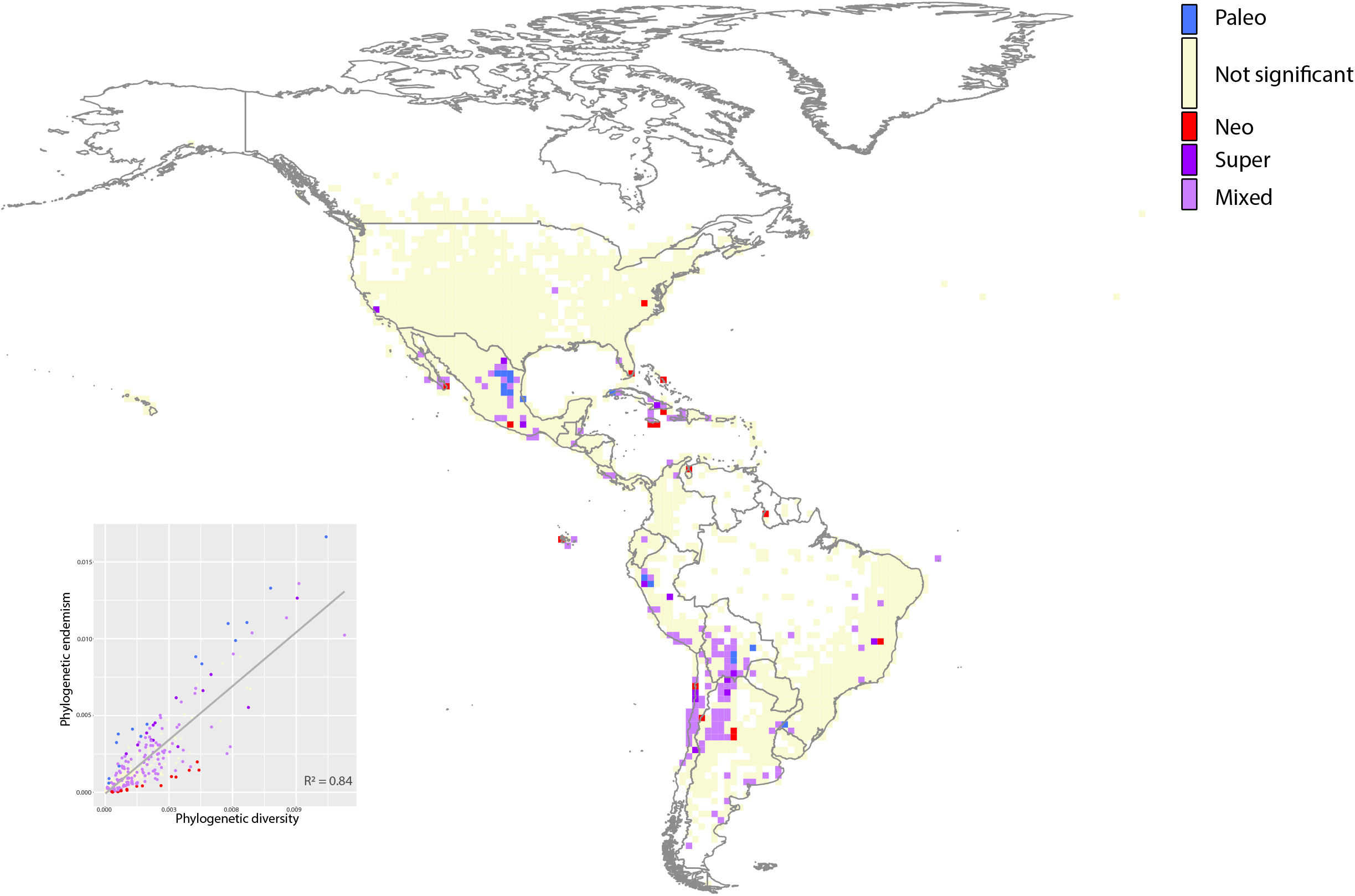
Spatial phylogenetics of Cactaceae showed areas of paleo-, neo-, and superendemism scattered in the Neotropical region. Red and blue cells show concentrations of significant neoendemism and paleoendemism, respectively; purple areas indicate concentrations of mixed endemism (both short and long branches present); dark purple indicates superendemism; beige cells show nonsignificant endemism areas. Scatter plots show the correlation between PD and PE.

### 3.3. Correlates of spatial diversity pattern

Past climatic, topography, and solar irradiance seem to highly contribute to the predictions of endemism, PD, and PE, while the soil features also displayed high contributions to the prediction of paleo-, neo-, mixed, and superendemism into the Cactaceae family (Fig. 3). The overall results of the four machine learning approaches used in this study show that the most influential factor on endemism and PE was isothermality (14.7% and 16.6%, respectively) during the Last Interglacial period (LIG), followed by terrain roughness (11.7% and 8.8%, respectively; Fig. 3a). The factor most predicting PD was direct normal irradiation (15.1%), followed by the mean temperature of the driest quarter of the past (11.1%; Fig. 3b). The soil texture and soil phases (15.9%) contributed highly to predict paleoendemism, followed by past isothermality (10.4%); to the neoendemism, the mean diurnal range (11.5%) and direct normal irradiation (11.2%) were the highly contributing predictors; to mixed endemism, past isothermality (16.4%), direct normal irradiation (12.1%), and terrain roughness (11.5%); and to superendemism, the highly contributing predictors were direct normal irradiation (12.4%), soil texture and soil phases (8.4%), and past isothermality (8.3%). Thus, the multimodel selection suggested a role of past temperature, soil, and solar irradiation as the main correlates of all kinds of endemism.

**Fig. 3.**
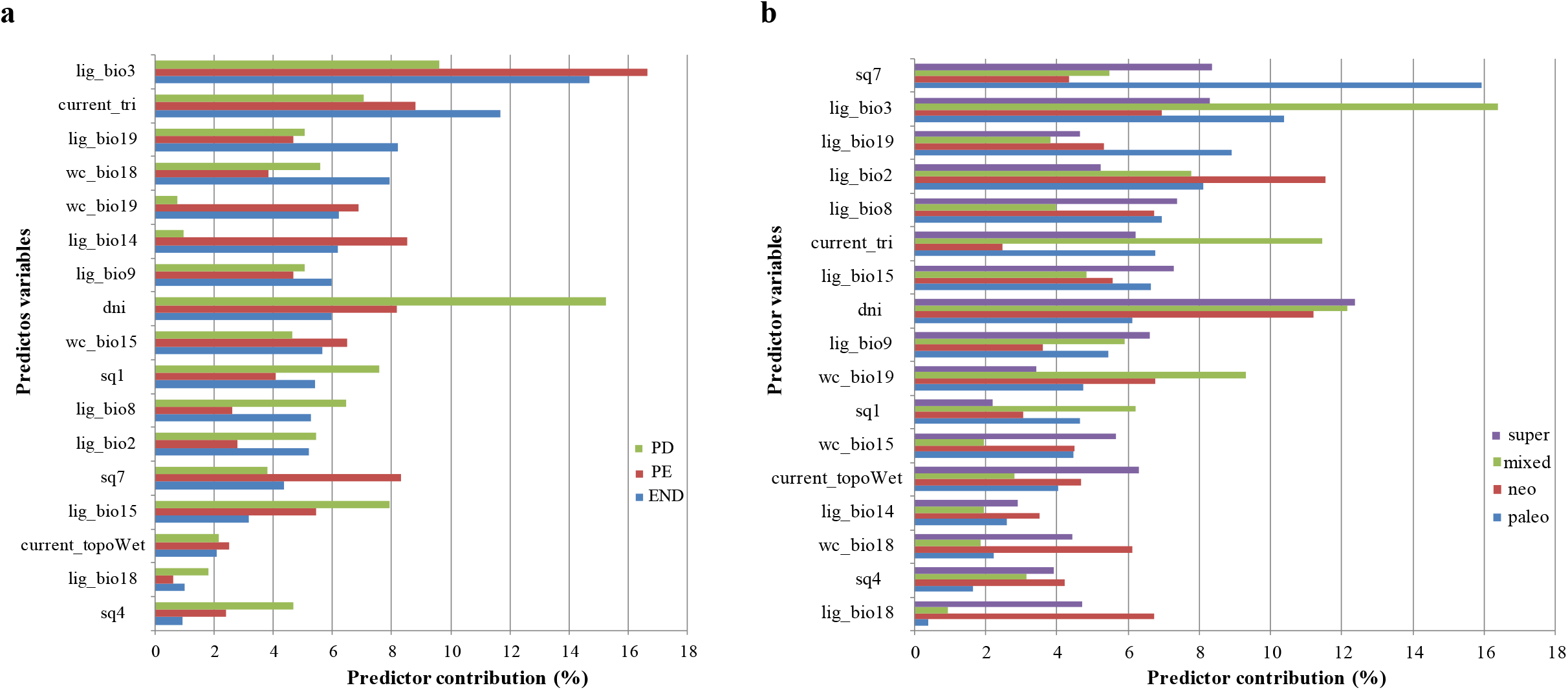
The relative importance (in percentage) of predictors of diversity for Cactaceae species. Panel (a) corresponds to predictor variables of the patterns of endemism, phylogenetic diversity (PD), and phylogenetic endemism, while in panel (b), we observed the predictors of the pattern of endemism diversity (see Table S1 for more details about variable description).

### 3.4. Protected areas associated with cacti diversity pattern hotspots

Here, we overlapped the grid cells of endemism, PD, PE, paleo-, neo-, and superendemism of cactus species to the legally protected areas present in the Nearctic and Neotropical regions (Fig. 4a-d). We showed that few cells associated with high diversity patterns were within legally protected areas (endemism cells: 13.8%; PD cells: 13.2%; PE cells: 16.25%; and superendemism: 28.8%). However, areas of neo- (52%) and paleo- (9.5%) seem to be better covered by these protected areas. Most of them are located in the United States, Mexico, and Central America. In South America, the few legally protected areas in Chile, which comprise a great part of the cactus endemism (including an area of super endemism; Fig. 4d), mainly in the Atacama Desert/Chilean Matorral, are evident. The Andes and the Campos Rupestres (Brazil) from Espinhaço Range were also poorly protected, even though both contained an enormity of endemic and microendemic species (Särkinen et al., 2012; Rapini et al., 2021). Thus, the analyses showed that only a small part of the cactus evolutionary history is legally protected (Goetsch et al., 2019; UNEP-WCMC and IUCN, 2021).

**Fig. 4.**
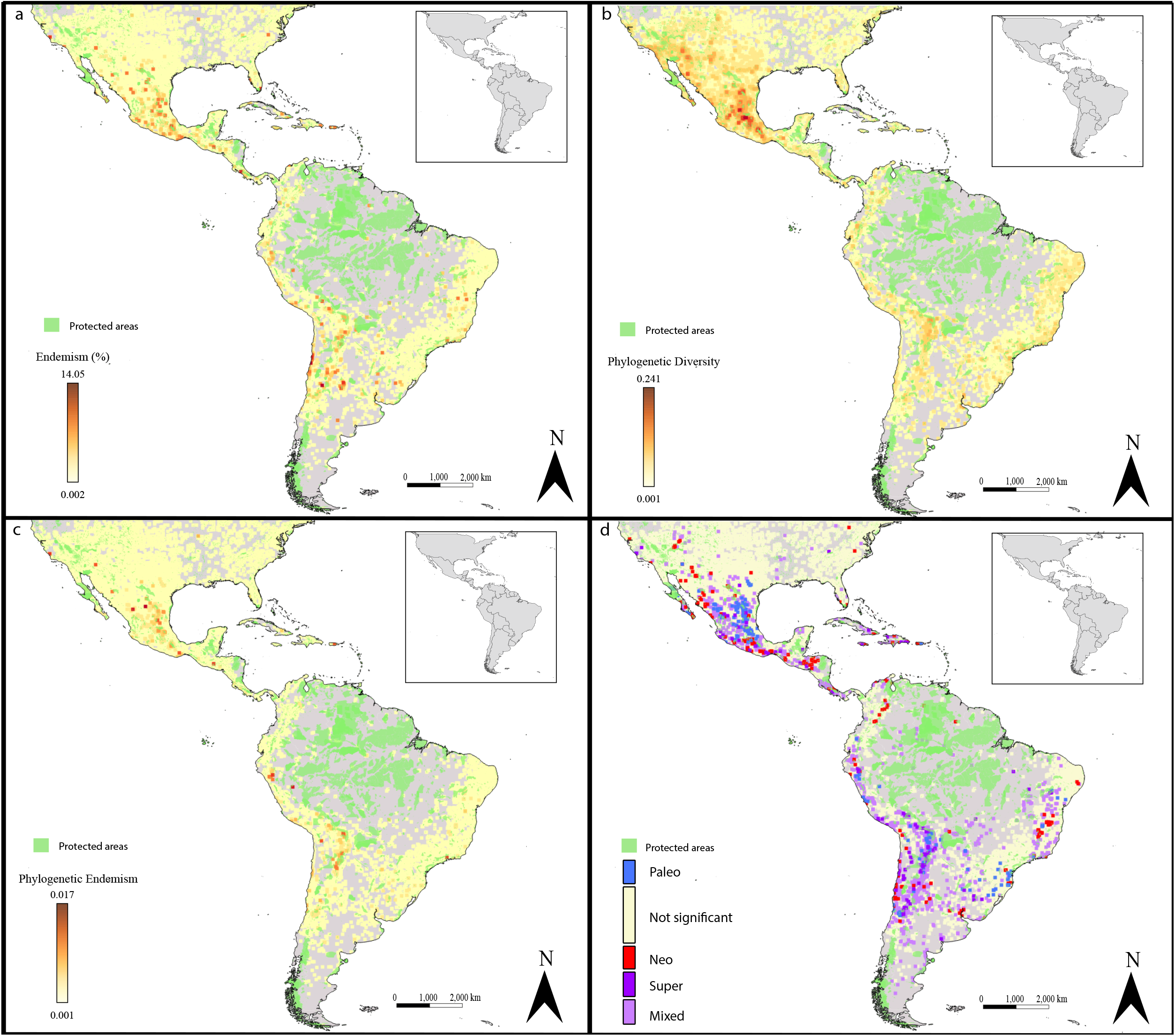
Endemism hotspot of Cactaceae with the legally protected areas of the Americas in green (UNEP-WCMC and IUCN, 2021). The map shows the low level of cells containing high (a) endemism, (b) phylogenetic diversity (PD), and (c) phylogenetic endemism (PE), and the possible areas of neo-, paleo-, and superendemism overlapped for the current formally protected areas.

## 4. Discussion

In this study, we integrated phylogenetic and spatial approaches to determine levels of diversity and endemism across the distribution of the family Cactaceae. Hotspots of cactus diversity were previously described (e.g. Rzedowski, 1993; Noroozi et al., 2018; Sosa et al., 2018; 2020; Nanni et al., 2019; Lavor et al., 2020), including deserts (Chihuahuan, Sonoran, and Atacama Deserts) and montane dry tropical forests (e.g., the Sierra Madre Oriental). However, these descriptions do not reflect the total complexity of endemism levels for the family. By adding empirical data based on different endemism metrics, we observed a higher diversity pattern also associated with the Atacama Desert, montane dry sub- and tropical forest (Chiquitania, Bolivian Montane Dry Forest, Chilean Matorral, Dry Chaco, etc.), patches of xeric vegetation within the Brazilian Atlantic Forest, and portions of the Espinhaço Range. It is worth noting that many of these areas are climatically stable (Hartley et al., 2015; Mucina and Wardell-Johnson, 2011; Werneck et al., 2012; Costa et al., 2018; Pie et al., 2018), and thought of as museums and cradles for drought-adapted plants (Murphy et al., 2015). Moreover, many areas of high PE are associated with the orogenic processes of the Miocene, such as the formation of the Gulf of California, the uplift of Mexican and Central American mountain systems, the Andes, and the Espinhaço Range (which extend from the Precambrian to the Cenozoic), which alter the local physiography, geological context, and drainage system (Saadi, 1995; Morán-Zenteno et al., 1999; De-Nova et al., 2018; Rech et al., 2019). The new geomorphological organization and topography may trigger local events of diversification and speciation, as observed in other studies (Steinbauer et al., 2016; Rull et al., 2020).

The significantly low values of PD recovered in areas of high endemism, such as the Sonoran and Atacama Deserts and a great part of the Caatinga and Atlantic Forests, may suggest strong lineage clustering (neoendemism), whereas the events of rapid and recent diversification of Cactaceae (Arakaki et al., 2011), such as the genus *Cereus* (Bombonato et al., 2020; Amaral et al., 2021), genus *Pilosocereus* (Bonatelli et al., 2014; Lavor et al., 2019), and Mammilloid (Breslin et al., 2021), favored the prevalence of closely related taxa. The significantly high PD in some areas, such as the Chihuahuan Desert, Peru and Ecuador Pacific coasts, the Andes, and the central portion of Dry Chaco, suggests these areas as historical refugia to the family and centers of diversification.

Most of the high PE areas comprise both ancient and young diversity (mixed endemism). However, areas of paleoendemism (museums) are encrusted within mixed-endemism cells (Fig. 2), suggesting a complex pattern of spatial endemism compartmentalization in some regions. These areas include the northern and central portions of the Andes and Atacama Desert/Matorral. At the same time, we observed neoendemism cells (possible cradles) in the peripheral regions with significantly high PE values (Fig. 3), which may indicate new centers of diversity.

Based on our diversity pattern results, deserts and drylands/dry forests may have acted as both museums and cradles of cactus lineages. This likely holds for other plants and animal groups in these landscapes (Sosa et al., 2018; 2020; Dick and Pennington, 2019; Candela et al., 2021). In the Neotropics, mainly in the Campos Rupestres from the Espinhaço Range, a major effort is underway to understand the environmental and evolutionary forces driving the increased rates of species richness and endemism in the old, climatically buffered, infertile landscapes (OCBILs; Hopper et al., 2021). Here, we found that the Campos Rupestres in the Espinhaço Range presents a high level of endemism and richness for Cactaceae (Fig. 1a-b). Indeed, similar results were reported for other species groups (Colli-Silva et al., 2019; Assunção-Silva et al., 2021). Here, we stress the Campos Rupestres from the Espinhaço Range as a center of superendemism to the South American cacti, being a remarkable hotspot of biodiversity and focus of conservation.

### 4.1. Predictor variables of Cactaceae endemism

The association between long-term climate stability and endemism is widely discussed in biogeographic and ecologic studies (Barrat et al., 2017; Feng et al., 2019; Zuologa et al., 2019). However, specific abiotic features may be more meaningful than others to explain differences in diversity patterns among species and/or bioregions (Barrat et al., 2017; Costa et al., 2018; Paz et al., 2020). Our results suggested that abiotic variables associated with temperature, soil roughness (topography), soil texture/phase, and solar irradiation are important factors that may explain the diversity pattern in Cactaceae.

The role of temperature and precipitation as drivers of diversification and bioregionalization has long been recognized (e.g., Antonelli, 2017). For Cactaceae, temperature seems to be the most important factor to predict the species diversity in the Neotropics, which was also demonstrated for specific genera of cactus and other groups of species distributed in xeric landscapes (López, 2017; Gottlieb et al., 2019; Mosco, 2019; Aquino et al., 2021). The isothermality (diurnal range of temperatures per temperature seasonality) in arid and semiarid regions, including desertic areas, displays low oscillations between day and night, which seems to favor CAM photosynthetic metabolism and the reduction in evapotranspiration of Cactaceae (stomata remaining closed during the daytime; Mosco, 2019). Thus, the arid areas may work as locally buffered zones over time, which might also have increased the diversity pattern into Cactaceae (Seal et al., 2017).

Landscape metrics, such as topographic roughness, revealed some endemism patterns in South America (Moeslund et al., 2013; Paz et al., 2020). We found that topography (terrain roughness) may act as a good predictor of endemism patterns in the Cactaceae family. The topographic heterogeneity associated with areas of highest PE values in our analysis, such as the Chihuahuan and Atacama Deserts and Andes, may produce a direct effect on cactus diversification. Guerrero et al. (2011) showed a complex pattern of diversification in the Atacama Desert, in which the topography offers several distinct habitats, also acting as a buffered area against climatic changes (Moeslund et al., 2013). Local soil patterns (e.g., soil pH and salinity) and texture are also important features directly affected by topography and act as predictors of species richness (Cingolani et al., 2010), which might represent an important influence on plant diversity, including desert plants (Moeslund et al., 2013; Muenchow et al., 2013). In the Cactaceae, the soil texture, such as that found in limestone outcrops, was demonstrated to be important to species persistence and, consequently, to diversification (Ruedas et al., 2006; Bárcenas-Argüello et al., 2010). For instance, Flores et al. (2019) show that the rock landform surface, which is a common substrate for many cactus species, is associated with species richness in arid environments. Thus, topographic roughness and soil texture may be proxies for habitat heterogeneity and diversification in arid and xeric regions. It is likely that other soil characteristics, such as pH and salinity, might be important predictors of diversity. Recently, Aquino et al. (2021) used a detailed Mexican soil database to highlight the relevance of soil pH in explaining patterns of distribution of the cactus genus *Epithelantha*.

As already demonstrated, cactus species are well adapted to grow and persist in environments with extreme temperatures and high solar radiation (Mauseth and Plemons-Rodriguez, 1999; Aliscioni et al., 2021), displaying particular morphological and physiological features to explore these habitats (Ehleringer et al., 1980; Albanese et al., 2019). Spines, cladode structure and shape, number of columns, and growth orientation are important features for minimizing absorption the solar radiation (Zavala-Hurtado et al., 1998; Menezes et al., 2015; Aliscioni et al., 2021), reducing damage processes associated with heat and light excess and balancing the amount of daily quanta absorbed into the photosynthetic system (Albanese et al., 2019). Recently, our research group showed positive selection signatures in genes associated with the photosynthetic system in *Cereus fernambucensis* (Cereeae), which reinforces the importance of solar irradiance within Cactaceae(Amaral et al., 2021). Furthermore, irradiance seems to influence flowering and fruiting evolution since seasonal variations in irradiance limits alter reproductive phenologies and seed development (Zimmerman et al., 2007). The importance of solar irradiance to cactus species may explain the function of this predictor in diversification and endemism in the family.

### 4.2. Conservation phylogenetics

We highlighted here areas with elevated diversity and endemism patterns for the Cactaceae family, including regions of broad interest to conservation. Approximately 30% of the Cactaceae species are currently considered critically endangered by the IUCN red list and should be threatened with extinction (Goettsch et al., 2015; 2019). Endangerous cactus species are of heightened concern due to their great scientific value and their significant social and economic role (Pedrosa et al., 2020; Tremlett et al., 2021). We identified that a great part of the cactus cell grids that comprise PD, PE, paleo-, neo-, and superendesmism regions are still out from legally protected areas (Fig. 4). These results indicate several unprotected areas of cactus diversity, some of which have already been proposed by Goettsch and collaborators (2015; 2019) using traditional diversity metrics, such as species richness and endemism. Here, we focus on the diversity pattern level, which encompasses ecological and evolutionary processes, showing that several of these areas are likely underestimated in extent and unrecognized as a protective unit. Despite the United States, Mexico, and some Central American countries displaying several protected areas that include arid and semiarid land (Fig. 4), the number of protected areas in South America is still insufficient.

The use of nontraditional metrics to estimate diversity patterns seems to provide new and additional information about biodiversity aspects, including evolutionary history associated with distinct geographic areas. This approach may help to improve conservation decisions. For instance, high-diversity cells that suggest rapid radiation events seem to be common among Cactaceae and may carry genetic variation potentially able to respond to future climate changes (Xu et al., 2019). Thus, cells that comprise paleo, neo-, and superendemism areas display high importance to the planning and management of strict protective units. Overall, less than 50% of the areas that we recovered in the analysis as of high conservation interest are unprotected. We also reaffirm the importance of species richness and endemic metrics, instead of disagreement in relation to their use. We proposed, like many other authors (Lee and Mishler, 2014; Laity et al., 2015; Xu et al., 2019), the incorporation of these metrics to improve the monitoring and planning of conservation areas.

Given the rapid anthropogenic disturbance, mainly in the past three decades, most of the areas outside and, even, inside the legally protected units have been lost due to predatory agriculture and cattle ranching and criminous fires, such as those observed in Brazil in the recent years (Pivello et al., 2021). Based on our results, distinct efforts may be established to plan protection areas, mitigate resource exploration, and manage threatened species in semiarid and arid lands. We propose to reinforce the conservative efforts to the maintenance of the already existing protected areas, mainly those that may recognize PE areas, not only for Cactaceae but also for all fauna and flora diversity. Investment in a better system/model to monitor biodiversity and the environment, rigorous and straightforward legally governmental politicians, and prioritization of regions of high biodiversity are fundamental for this purpose. Furthermore, the planning and establishment of new protected units, mainly in South America, using diversity metrics may improve and upgrade the preservation of the management board.

### 4.3. Caveats

Here, we propose a first attempt to cover the general diversity patterns related to the cactus species richness distributed in the Neotropical xeric landscapes. It is expected that by using high-throughput sequencing technologies, more complete datasets might be available in the future (including new taxa), optimizing the branch length estimates and, consequently, the metrics reported here. New and improved public databases that include specific variables with an increased resolution (e.g. soil pH) may also promote more detailed cell grids with diverse endemic patterns. Thus, we seek to standardize the metrics of analyses and genetic and occurrence sampling to minimize these caveats, conscious of the barriers and biases associated with this approach. In spite of this, we were able to define areas of ecological importance to the maintenance and diversification of the family, displaying the main areas deserving conservatism efforts and describing the importance of protective unit planning to the Neotropics.

## Acknowledgment

We thank the Fundação de Amparo à Pesquisa do Estado de São Paulo (FAPESP) for two research grants (2018/03428-5 to F.F.F. and E.M.M. 2021/15161-3 to F.F.F.) and a fellowship (2018/06937-8 to M.R.B). This study was also financed in part by the Coordenação de Aperfeiçoamento de Pessoal de Nível Superior - Brasil (CAPES) - Finance Code 001 (fellowship to D.T.A.). E.M.M. thanks National Council for Scientific and Technological Development (CNPq, grant 03940/2019-0) for financial support.

## Data Accessibility Statement

The dataset supermatrices of 40%, 60%, and 80% missing data, as well as the tree topology obtained for the three datasets, were deposited on FigShare (10.6084/m9.figshare.17057816).

