## Supplementry material for "Spatial patterns of evolutionary diversity in Cactaceae show low ecological representation within protected areas"

**Appendix A. Glossary**

**Taxon richness (TR):** Count the number of species in the neighbor sets

**Weighted endemism (WE):** Calculate endemism for labels inversely weighted by species ranges

**Phylogenetic diversity (PD):** a measure of biodiversity that depends on the tree topology. Calculate a set of species as equal to the sum of the lengths of all those branches on the tree that span the members of the set.

**Phylogenetic endemism (PE):** a spatial measure of relative phylogenetic diversity in an assemblage weighted by range size, measured as the sum of branch length of descendant species in an assemblage weighted by the descendant range sizes.

**Relative phylogenetic diversity (RPD)**: The ratio of the tree's PD to a null model of PD evenly distributed across all nodes. This metric discerns any correlations or patterns in the data layers over the randomization maps of RPD.

**Relative phylogenetic endemism (RPE)**: The ratio of the tree's PE to a null model where PE is calculated using a tree where all branches are of equal length. This metric discerns any correlations or patterns in the data layers over the randomization maps of RPE.

**Paleoendemism**: species that were formerly widespread but are now restricted to reduced area distribution**.**

**Neoendemism**: the ecological state of a species being unique to a defined geographic location, which they have recently arisen.

**Mixed-endemism:** cells that both paleo- and neoendemism are observed.

**Super-endemism:** extremely high levels (> 0.97) of both neo- and palaeo-endemism.

**Appendix B.** Questions and hypotheses table associated with this study

**Question:** Which are those areas with high phylogenetic endemism (PE) and diversity (PD) within Cactaceae distribution? Are they associated with the diversification of Cactaceae? Where are the neo- and paleo- endemism hotspots for these species? Which abiotic factor may explain these PD and PE patterns? Which areas are of conservation interest?

**Hypotheses tested in this work:**

| Hypotheses | Mechanism | Prediction | Analyses | Acceptance |
|---|---|---|---|---|
| Patterns of species richness and lineage diversification in Cactaceae | Areas with high PD and PE values, which comprises the SR metrics, have been stable for long time periods | High PD and PE values in specific patches | Phylogenetic Diversity metrics (Biodiverse) | Yes |
| Areas with higher PD and RPD values in Cactaceae, such as rocky outcrops, highlands, and arid and xeric environments are considered biodiversity hotspots to this group | Long stable xeric areas may accumulate more lineages over time, long-surviving species, and older lineages. | Clades with long branches and structured clades | Species Richness measure and Phylogenetic Diversity (Biodiverse 3.1); Neo- and Paleo-endemism (Biodiverse and CANAPE) | Yes |
| Cactaceae endemism may work as cradles and museums of species | Neoendemism is more likely in current xeric and open areas, whereas areas of paleoendemism will be more likely where climate has been stable. | Higher values of PE and RPE in recent, older, or mixed areas | Neo- and Paleo-endemism (Biodiverse and CANAPE) | Partially |
| | | | Test the importance of abiotic drivers (machine learning approaches) | Yes |
| Areas with high PD, RPD, PE, and RPE should be focused on the conservation purposes | Neoendemism is more likely in climatically unstable and landscapes areas, which may be protected | Areas with higher values of PE and RPE are associated with suitable areas to the species | Phylodiversity metrics (Biodiverse and CANAPE) | Yes |

able S1. The compiled environmental variables used as diversity pattern predictors in this study. The variables marked with the asterisks show the seventeen variables used in the study fter excluding highly correlated variables.

| Variable | Description | Source | Category |
|---|---|---|---|
| Bio1 | Annual Mean Temperature | WorldClim | Current temperature |
| Bio2 | Mean Diurnal Range | | |
| Bio3 | Isothermality (BIO2/BIO7) (* 100) | | |
| Bio4 | Temperature Seasonality (standard deviation *100) | | |
| Bio5 | Max Temperature of Warmest Month | | |
| Bio6 | Min Temperature of Coldest Month | | |
| Bio7 | Temperature Annual Range (BIO5-BIO6) | | |
| Bio8 | Mean Temperature of Wettest Quarter | | |
| Bio9 | Mean Temperature of Driest Quarter | | |
| Bio10 | Mean Temperature of Warmest Quarter | | |
| Bio11 | Mean Temperature of Coldest Quarter | | |
| Bio12 | Annual Precipitation | WorldClim | Current precipitation |
| Bio13 | Precipitation of Wettest Month | | |
| Bio14 | Precipitation of Driest Month | | |
| Bio15* | Precipitation Seasonality (Coefficient of Variation) | | |
| Bio16 | Precipitation of Wettest Quarter | | |
| Bio17 | Precipitation of Driest Quarter | | |
| Bio18* | Precipitation of Warmest Quarter | | |
| Bio19* | Precipitation of Coldest Quarter | | |
| LIG_Bio1 | Annual Mean Temperature | PaleoClim | Last Intergratial Temperature |
| LIG_Bio2* | Mean Diurnal Range | | |
| LIG_Bio3* | Isothermality (BIO2/BIO7) (* 100) | | |
| LIG_Bio4 | Temperature Seasonality (standard deviation *100) | | |

| | | | |
|---|---|---|---|
| LIG_Bio5 | Max Temperature of Warmest Month | | |
| LIG_Bio6 | Min Temperature of Coldest Month | | |
| LIG_Bio7 | Temperature Annual Range (BIO5-BIO6) | | |
| LIG_Bio8* | Mean Temperature of Wettest Quarter | | |
| LIG_Bio9* | Mean Temperature of Driest Quarter | | |
| LIG_Bio10 | Mean Temperature of Warmest Quarter | | |
| LIG_Bio11 | Mean Temperature of Coldest Quarter | | |
| LIG_Bio12 | Annual Precipitation | PaleoClim | Last Intergratial Precipitation |
| LIG_Bio13 | Precipitation of Wettest Month | | |
| LIG_Bio14* | Precipitation of Driest Month | | |
| LIG_Bio15* | Precipitation Seasonality (Coefficient of Variation) | | |
| LIG_Bio16 | Precipitation of Wettest Quarter | | |
| LIG_Bio17 | Precipitation of Driest Quarter | | |
| LIG_Bio18* | Precipitation of Warmest Quarter | | |
| LIG_Bio19* | Precipitation of Coldest Quarter | | |
| topoWET | SAGA-GIS topographic wetness index | Envirem | Topography |
| TRI* | terrain roughness index | | |
| Sq1* | Nutrient availability | Harmonized World Soil Database | Soil |
| Sq2 | Nutrient retention capacity | | |
| Sq3 | Rooting conditions | | |
| Sq4* | Oxygen availability to roots | | |
| Sq5 | Excess salts | | |
| Sq6 | Toxicity | | |
| Sq7* | Workability (Soil texture and soil phase) | | |
| DNI* | Direct Normal Irradiation | Solargis | Solar irradiation |
| DHI | Direct Horizontal Irradiation | | |

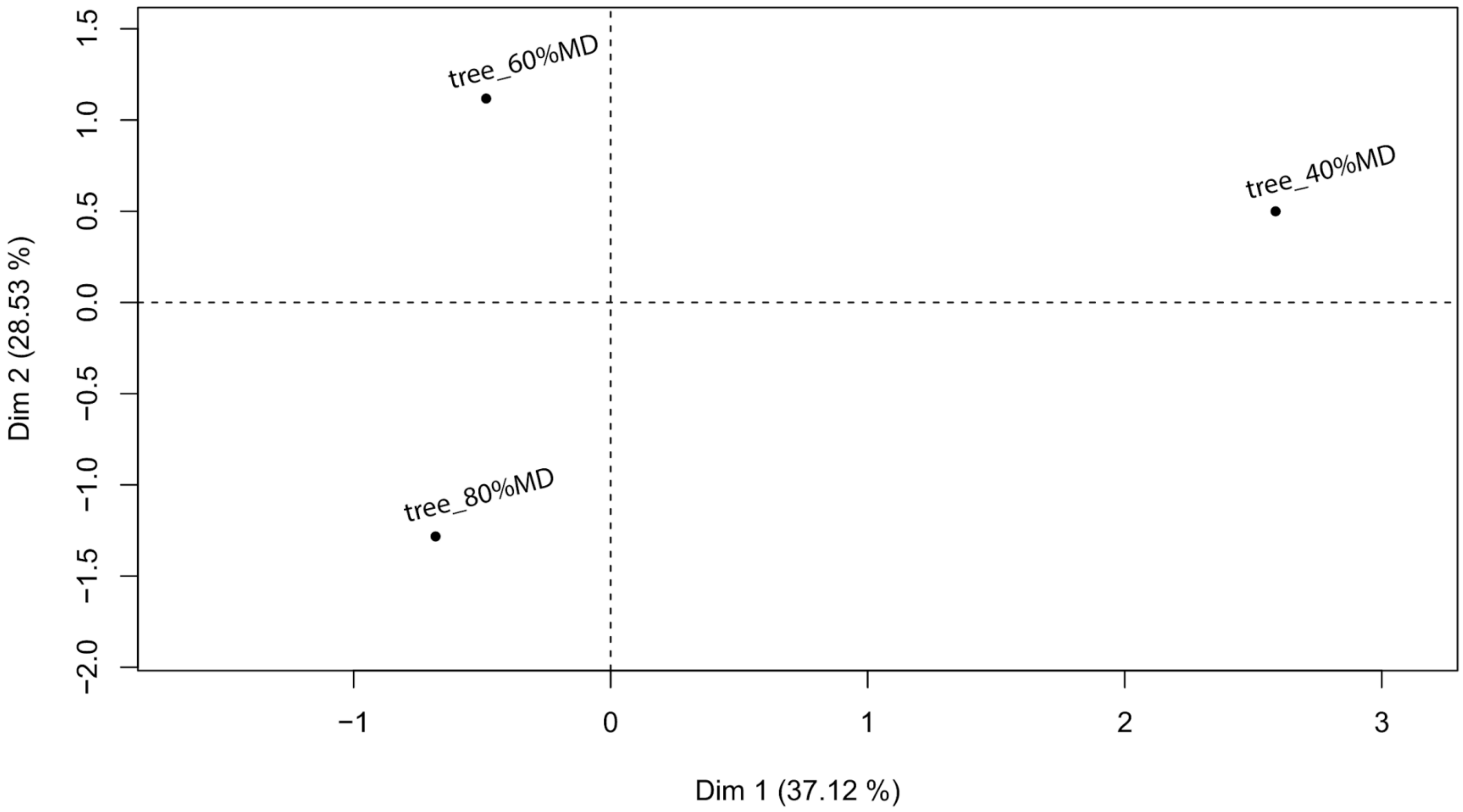


**Figure S1.** PCA based on Robinson–Foulds (RF) distances between the distinct maximum likelihood topologies generated by distinct datasets (40%a, 60%, and 80% of missing data).

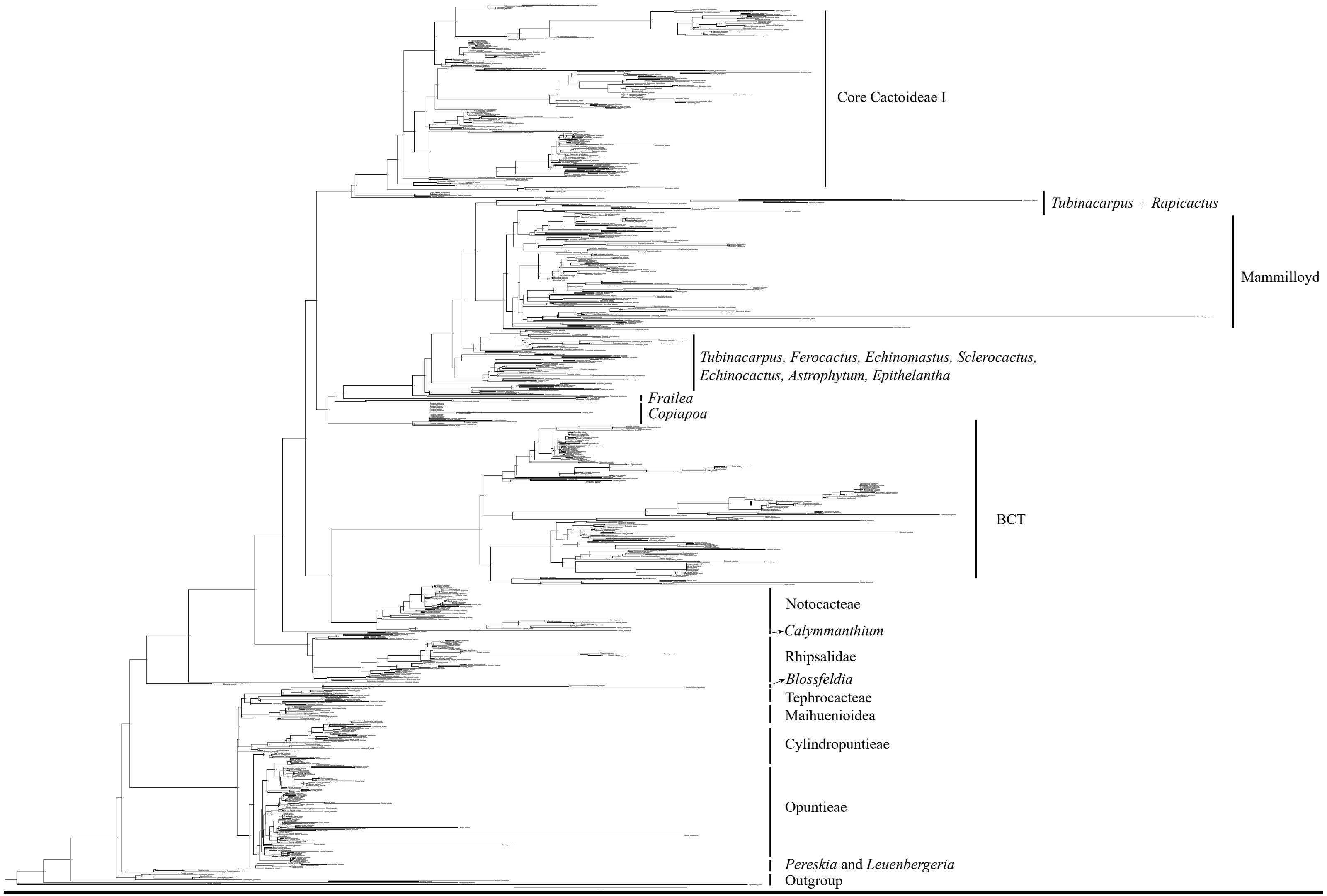


**Figure S2.** The tree topology used in this study. The clades were classified as proposed by Guerrero et al. (2019).
